# Re-infection with SARS-CoV-2 is associated with increased antibody breadth and potency against diverse sarbecovirus strains

**DOI:** 10.1101/2025.10.03.679858

**Authors:** Michelle Lilly, Felicitas Ruiz, Will Foreman, Vrasha Chohan, Jamie Guenthoer, Delphine Depierreux, Viren A. Baharani, Duncan Ralph, Alex Harteloo, Helen Y. Chu, Paul D. Bieniasz, Tyler N. Starr, Julie Overbaugh

## Abstract

The ease with which emerging SARS-CoV-2 variants escape neutralizing antibodies limits protection afforded by a prior exposure, be it infection or vaccination. While rare, broadly neutralizing antibodies with activity towards diverse sarbecoviruses have been detected in convalescent serum. Motivated by findings that plasma responses show increased neutralization breadth and potency with continued antigen exposure, we isolated monoclonal antibodies (mAbs) after a SARS-CoV-2 re-infection and compared them to those isolated one year prior, after the first breakthrough infection. Among clonal lineage members identified at both time points, mAbs from the later time point showed improved neutralization potency and breadth. One mAb isolated after re-infection, C68.490, targets a conserved region in the receptor binding domain core and shows remarkable activity not only against SARS-CoV-2 variants, but also diverse sarbecoviruses from more distant clades present in animal reservoirs. These findings suggest that a focus on individuals with diverse and repeated antigen exposure could lead to identification of antibodies with therapeutic utility not just towards current and future SARS-CoV-2 variants, but also distant sarbecoviruses in the event of a future spillover.

## INTRODUCTION

The SARS-CoV-2 pandemic has illustrated the devastating effects that can occur when a sarbecovirus originally from an animal reservoir is introduced into the human population. From this experience, we have come to appreciate the important clinical contribution of neutralizing antibodies to protection against sarbecovirus infections and disease severity. Neutralizing antibodies contribute to SARS-CoV-2 vaccine efficacy and are a correlate of protection from infection and disease (Goldblatt et al. 2022). In addition, the therapeutic use of monoclonal antibodies (mAbs) proved effective in decreasing disease severity due to early strains of SARS-CoV-2 infection before viral escape mutations arose (Chen Peter et al. 2021; Copin et al. 2021; Gupta et al. 2021; Weinreich, Sivapalasingam, Norton, Ali, Gao, Bhore, Xiao, et al. 2021; Weinreich, Sivapalasingam, Norton, Ali, Gao, Bhore, Musser, et al. 2021). Neutralizing antibodies also showed promise against SARS-CoV-1, although they were not deployed due to the curtailment of that more limited sarbecovirus outbreak (ter Meulen et al. 2006; Wong et al. 2003). Characterizing neutralizing antibodies with broad sarbecovirus activity is therefore important for pandemic preparedness and prevention, given that sarbecoviruses are prevalent in a variety of animal reservoirs (Ge et al. 2013; Roelle et al. 2022; Seifert et al. 2022; Wacharapluesadee et al. 2021) and have recently seeded several human outbreaks.

Cross-reactive neutralizing antibodies that target functionally constrained epitopes across sarbecoviruses may also prove more durable against SARS-CoV-2 variation, which continues to compromise the efficacy of many neutralizing antibodies. Because the immune response generated by vaccination and infection is not sterilizing, the virus replicates for days, or sometimes weeks, in the face of neutralizing antibodies generated by these exposures. The most potent of these neutralizing antibodies typically target the receptor binding domain (RBD) in the S1 subunit of the viral spike glycoprotein (Dejnirattisai et al. 2021; Greaney et al. 2021; Piccoli et al. 2020; Qi et al. 2022; Robbiani et al. 2020). Not surprisingly then, the antibody escape observed for SARS-CoV-2 variants is driven mostly by mutations in dominant RBD epitopes, which show remarkable ability to tolerate variation while retaining function (Cao et al. 2023; Starr et al. 2020; Taylor and Starr 2024; Witte et al. 2023). Unfortunately, this escape has occurred in epitopes targeted by therapeutic mAbs, rendering them ineffective and highlighting a need for mAbs that target more mutation-intolerant regions of the RBD (Cameroni et al. 2022; Cao et al. 2023; Cao, Wang, et al. 2022; Cao, Yisimayi, et al. 2022).

Sequence variation within RBD is not uniform and some regions show very limited variation across circulating strains, suggesting there may be functionally constrained regions in the spike RBD. Indeed, several RBD-specific mAbs targeting more conserved regions that exhibit broad sarbecovirus neutralizing activity have been identified (Cameroni et al. 2022; Tortorici et al. 2021; Cao, Jian, et al. 2022; Starr et al. 2021; Cui et al. 2024; Park et al. 2022; Rappazzo et al. 2021; Jensen et al. 2023; Pinto et al. 2020; Jette et al. 2021; Liu et al. 2020; K.-Y. A. Huang et al. 2023; Guenthoer et al. 2023; Ruiz et al. 2024). S2X259 was one of the first pan-sarbecovirus neutralizing antibodies characterized, although it rapidly lost activity against emerging SARS-CoV-2 Omicron variants (Rosen et al. 2024; Ruiz et al. 2024; Cao, Jian, et al. 2022). Subsequently, antibodies with impressive SARS-CoV-2 breadth have been isolated from individuals who were exposed to both SARS-CoV-1 and SARS-CoV-2. One notable example is SA55, which retains breadth and potency across all SARS-CoV-2 strains tested, including recent variants (Cao, Jian, et al. 2022; Planas et al. 2024). SA55 also exhibits binding and neutralizing capabilities against SARS-CoV-1 as well as a subset of sarbecoviruses from different clades (Cao, Jian, et al. 2022; Rosen et al. 2024). A recently described potently neutralizing and cross-reactive antibody, VIR-7229, was obtained by mimicking the affinity maturation process through selection for mutations that led to improved binding *in vitro* (Rosen et al. 2024). This mAb showed impressive binding breadth across all major SARS-CoV-2 variants and against all but a small subset of sarbecovirus spike proteins, although it exhibits more limited neutralization potency against some SARS-CoV-2 variants (Rosen et al. 2024). There are examples of mAbs that are broadly neutralizing against SARS-CoV-2 variants and show some cross-reactivity towards distant sarbecoviruses, and conversely, there are mAbs that show cross-sarbecovirus breadth but limited SARS-CoV-2 neutralization, but there are few mAbs that do both (Rappazzo et al. 2021; Cao, Jian, et al. 2022; Cameroni et al. 2022; K.-Y. A. Huang et al. 2023). Such antibodies, which would be predicted to target highly conserved and constrained epitopes, are likely to be relatively resistant to escape.

We previously identified several broadly neutralizing antibodies in one individual, C68, that experienced a heterologous antigen exposure to SARS-CoV-2 spike through a Delta breakthrough infection (BTI) subsequent to vaccination with the ancestral WH-1 strain (Guenthoer et al. 2023; Ruiz et al. 2024). One of these mAbs (C68.61) targets a conserved region in the core of the RBD where there has been minimal antigenic variation in SARS-CoV-2 variants (Guenthoer et al. 2023) and escape could not be selected *in vitro* (Ruiz et al. 2024). Interestingly, C68.61 shows breadth not only across all SARS-CoV-2 variants tested, but also against a subset of sarbecovirus variants from animal reservoirs (Ruiz et al. 2024).

Given the resilient functional properties of mAbs from C68, we sought to determine whether there might be mAbs with even greater breadth or potency after subsequent antigen exposures by vaccinations and infection to genetically diverse SARS-CoV-2 variants in this individual. Several studies have examined plasma responses and shown increases in breadth and potency of neutralization with repeated exposure to heterologous spike antigen in polyclonal sera (Wang et al. 2023, 2024; Abbad et al. 2024; Evans et al. 2022), although detailed studies of affinity maturation of individual monoclonal responses in the context of heterologous spike exposures are few (Kaku et al. 2023; Kotaki et al. 2024). Likewise, variation in SARS-CoV-2 may provide antigenic diversity that could drive the development of a more potent and/or broad neutralizing antibody response through somatic hypermutation (SHM) and affinity maturation. In line with this, individuals with both a WH-1 and an Omicron variant exposure developed higher and broader plasma neutralizing titers against Omicron variants than individuals with multiple exposure to the ancestral WH-1 strain by vaccination (Kurhade et al. 2023; Cao et al. 2023; Collier Ai-ris Y. et al. 2023; Quandt et al. 2022).

To examine the process of affinity maturation of SARS-CoV-2-binding antibodies and identify cross-reactive neutralizing antibodies, we isolated spike-specific B cells from a later time point in the individual from whom C68.61 and other broadly active mAbs were isolated. We selected a sample obtained after an additional vaccination and a second BTI, which occurred during a period when Omicron variants were the major circulating strains. We characterized the genetic and functional properties of mAbs reconstructed from B cells that bound spike antigen, focusing on antibodies that were part of clonal lineage families to examine affinity maturation. We observed increased neutralization breadth and potency against SARS-CoV-2 variants in clonal lineage members isolated from this later time point compared to those from the earlier sample. Remarkably, we also identified a mAb, C68.490, from a clonal lineage not detected in earlier timepoints that showed breadth and potency against SARS-CoV-2 variants as well as multiple sarbecoviruses representing several clades common in animal reservoirs.

## MATERIALS AND METHODS

### Participant information

A peripheral blood mononuclear cell (PBMC) sample collected from individual C68 in June 2022, one month after a second BTI, was used for this study. C68 had a first BTI with a Delta variant in July 2021, which was two months after immunization with two doses of the Pfizer-BioNTech mRNA vaccine. mAbs were previously isolated one month after this BTI [referred to here as the BTI-1 time point (Guenthoer et al. 2024)]. For the current study, the sample analyzed was one year later (BTI-2 time point), which was a month after a second infection, at a time when the Omicron BA.5 lineage was the dominant circulating variant, and 8 months after a third vaccination. All vaccinations were with the ancestral WH-1 strain. Study participant C68 was enrolled in the Hospitalized or Ambulatory Adults with Respiratory Viral Infections (HAARVI) study. This study was approved by University of Washington Protocol #STUDY00000959 with reliance from the institutional review board at the Fred Hutchinson Cancer Center.

### B cell isolation approach

The PBMC sample from BTI-2, which contained 11.2 million viable cells, was stained and sorted to isolate spike-specific memory B cells (MBCs) as previously described (Guenthoer et al. 2023). In brief, cells were washed in FACS wash and stained with a cocktail of fluorescently-labeled antibodies to cell surface markers: CD3-BV711 (BD Biosciences, clone UCHT1), CD14-BV711 (BD Biosciences, cloneMφP9), CD16-BV711 (BD Biosciences, clone 3G8), CD19-BV510 (BD Biosciences, clone SJ25C1), IgM-FITC (BD Biosciences, clone G20-127), IgD-FITC (BD Biosciences, clone IA6-2). PBMCs were also incubated with biotinylated SARS-CoV-2 Omicron XBB.1.5 spike trimer (Acro Biosystems, cat. SPN-C82Ez-25ug) and biotinylated SARS-CoV-1 spike trimer (Acro Biosystems, cat. SPN-S82E3-25ug; CUHK-W1 strain, accession #: AY278554.2). Biotinylated spike proteins were conjugated to streptavidin-fluorophores APC and PE (from BioLegend) overnight at 4°C the day before cell sorting. PBMCs were loaded onto a BD FACS Aria II cell sorter, and IgG expressing, spike-specific B cells were identified as CD3−, CD14−, CD16−, CD19+, IgD−, IgM−, PE+, APC+ and live cells (Ghost Dye Red 780; Tonbo Biosciences cat: 13-0865-T500) and were sorted into a total of four 96-well PCR plates containing cell lysis/RNA storage buffer and stored at −80C. A total of 384 SARS-CoV-2 Omicron XBB.1.5 spike trimer and/or SARS-CoV-1 spike trimer-specific B cells were recovered from this sample.

### Recombinant antibody production and purification

Amplification of variable heavy and light chain genes (VH and VL, respectively) were performed as described (Guenthoer et al. 2023; Tiller et al. 2008; J. Huang et al. 2013). In order to identify clonal lineage members, the 384 B cells that bound spike were first subject to PCR to amplify the VH gene fragment and the sequence of the resulting PCR products were determined.

A total of 269 VH gene sequences were successfully amplified, with 254 predicted to encode productive VH genes based on the presence of a variable, diversity and joining (VDJ) gene rearrangement with in-frame open reading frame and with no premature stop codons.

To identify the subset that were members of the same clonal lineage, clonality was assessed based on VDJ gene segment rearrangement using *partis*, a BCR sequence annotation software package designed for antibody sequence analysis (Ralph & Matsen, 2016, 2022). This analysis was performed for the 269 BTI-2 VH sequences, of which 254 were predicted to encode productive genes, combined with 118 BTI-1 VH sequences from prior studies, which included 100 VH genes that were further functionally characterized (Guenthoer et al. 2023; Ruiz et al. 2024; Guenthoer et al., 2024). When clonal lineages were identified based on the VH sequence, either within the BTI-2 sequences or between BTI-2 and BTI-1 sequences, the VL chain sequence was amplified from select MBCs. Specifically, as a means to focus on those with known neutralizing activity, in cases where BTI-2 antibodies were clonal to a BTI-1 antibody, the lineages where the BTI-1 antibody was RBD-specific were pursued further. In cases where there were clonally related VH sequences within BTI-2 but no BTI-1 clonal counterparts, we rescued the VL gene corresponding to the antibody sequence with the highest SHM in the VH gene as defined by *partis* (Ralph & Matsen, 2016, 2022). In cases where the antibody with the highest SHM could not be isolated or cloned, another antibody from the same clonal lineage was selected, if available.

After the isolation of VL genes of interest, sequences were again assessed for clonality using *partis* in order to improve clonal lineage determinations with the addition of paired heavy-light chain sequences (Ralph and Matsen 2022). This second determination based on both VH and VL sequences was used in the final analysis. Based on the secondary *partis* determination, a total of 33 clonal lineages were identified, of which 22 had lineage family members within BTI-2 only and 11 had lineage family members in both BTI-2 and BTI-1.

mAbs were generated by transfecting FreeStyle 293-F Cells (Invitrogen) with paired VH and VL chains as described (Guenthoer et al. 2023). The resulting mabs were purified by affinity chromatography using protein G agarose columns (Pierce).

### ELISA binding assay

Antibody binding to SARS-CoV-2 or SARS-CoV-1 recombinant spike protein was assessed using ELISA binding assays as previously described (Garrett et al. 2021). All antibodies were tested at a single concentration (1000 ng/ml), and OD450 nm values were measured. The raw OD450 values were averaged between technical duplicates and background subtracted with the negative control wells treated with HIV mAb VRC01. All described antibodies were tested for binding to SARS-CoV-2 WH-1 RBD (Sino Biological, cat. 40592-V08H), SARS-CoV-2 WH-1 S1 (Sino Biological, cat. 40591-V08H), SARS-CoV-2 XBB.1.5 trimer (40589-V08H45), and SARS-CoV-1 GD01 trimer (Acro Biosystems, cat. SPN-S52Ht). In addition, a subset was also tested for binding to SARS-CoV-2 Delta trimer (Sino Biological, cat. 40589-V08H10), and SARS-CoV-2 B.Q.1.1 trimer (Sino Biological, cat. 40589-V08H41).

### Neutralization assay

Antibodies were tested for neutralization using spike-pseudotyped lentiviral particles and 293T-ACE2 expressing cells (Crawford et al., 2020) in a 384-well format, as described previously (Guenthoer et al. 2023). Sarbecovirus spike plasmids with spike genes specific for SARS-CoV-2 variants, SARS-CoV-1 (Urbani strain), BtKY72 (K493Y/T498W mutant), Khosta-2, WIV1, LYRa3, Pangolin-GD, RsSHC014 were generated as previously described (Guenthoer et al. 2024, 2023; Ruiz et al. 2024). Antibody dilutions were prepared at a starting concentration of 20 μg/mL, with three-fold dilution in a total of six dilutions assessed, with the exception of C68.490, and control antibodies C68.61 and S2X259, where two-fold mAb dilution curves for a total of 12 dilutions were used. The inhibitory concentration at 50% (IC50) was calculated with GraphPad PRISM statistical software by fitting a four-parameter (agonist vs response) nonlinear regression curve with the bottom fixed at 0, the top constrained to 1 and HillSlope < 0. In cases where fraction infectivity did not cross 0.5, the experiment was repeated with a lower starting mAb concentration so that the neutralization profile encompassed a sigmoidal curve with data points above 50% infectivity.

### Antibody binding to yeast-displayed pan-sarbecoviruses RBD

To evaluate the ability of C68.490 to bind a panel of sarbecovirus RBDs, we used yeast libraries displaying RBDs from sarbecoviruses representing different clades as previously described (Rosen et al. 2024). A full list of all RBDs and sequence accession numbers is available https://github.com/tstarrlab/SARSr-CoV_mAb-breadth_Overbaugh/blob/main/data/sarbeco_accessions.csv. Each sarbecovirus RBD is represented by multiple (often >100) barcodes such that technical pseudo-replicates can be ascertained within each binding experiment. C68.490 was incubated with a sarbecovirus-RBD-displaying yeast library at five concentration points (10,000 ng/mL and four serial 25-fold dilutions to 0.0256 ng/mL; plus zero mAb condition). Cells displaying RBDs were sorted via FACS into 1 of 4 bins according to binding towards C68.490. To set these bins, a known-binding pair of isogenic yeast-displayed RBD and antibody were incubated at saturation (10,000 ng/mL) and 0 ng/mL (no antibody). Bins were drawn such that bin 1 captured 95% of cells in the 0 ng/mL population, and bin 4 captured 95% of cells in the 10,000 ng/mL population. Barcodes were counted within each FACS bin via Illumina sequencing, with raw barcode counts available at https://github.com/tstarrlab/SARSr-CoV_mAb-breadth_Overbaugh/blob/main/results/counts/variant_counts.csv. EC50s were calculated by fitting the Hill equation (where n=1) to bin values (1-4) for each barcoded sarbecovirus RBD across the antibody concentration series, allowing us to compare an antibody’s affinity towards sarbecovirus RBDs in the library. EC50 scores are the geometric mean across the independent barcodes. The computational pipeline for computing sarbecovirus mAb-binding breadth is available from https://github.com/tstarrlab/SARSr-CoV_mAb-breadth_Overbaugh.

### Epitope profiling by deep mutational scanning of the RBD

To determine the epitope targets and the escape profile of the C68.490 mAb, we used a yeast library displaying RBD proteins with nearly every single amino acid mutation in the RBD of SARS-CoV-2 Wuhan-Hu-1, SARS-CoV-2 Omicron BA.2, and SARS-CoV-1 Urbani as previously described (Greaney et al. 2021; Guenthoer et al. 2023; Ruiz et al. 2024). A FACS-based approach was used to identify antibody escape mutants by gating on yeast that display RBD mutants based on antibody binding, with gates drawn on wildtype control cells labeled at 10% of the library selection mAb concentration to approximate an escape bin of 10x or greater loss of binding (Supplemental Fig 1A), sorted on a BD FACS AriaII.

Pre-sort and sorted antibody-escape cells were sequenced using an Illumina NovaSeq and a 16-nucleotide barcode linked to each RBD mutant was used to identify the mutant sequence. Raw sequence read counts are available from GitHub: https://github.com/tstarrlab/SARS-CoV-2-RBD_Omicron_MAP_Overbaugh-C68-490/tree/main/results/counts. Sequencing reads were then compiled and compared to the pre-sort population frequencies to generate the “escape fraction” visualized on logoplots (Supplemental Fig 1B). Experiments were performed in biological duplicate using independent mutant RBD libraries that correlate well (Supplemental Fig 1C) so escape fractions represent the average of these two independent biological replicates. Final escape fraction measurements averaged across two replicates are available from https://github.com/tstarrlab/SARS-CoV-2-RBD_Omicron_MAP_Overbaugh-C68-490/tree/main/results/supp_data. The entire pipeline for epitope escape profiling is available from https://github.com/tstarrlab/SARS-CoV-2-RBD_Omicron_MAP_Overbaugh-C68-490.

### Escape mutations identified by replication competent recombinant virus surface expressed spike protein

To assess which escape mutants arise in the presence of C68.490 mAb in the context of virus replication, we used a recombinant replication-competent vesicular stomatitis virus (rVSV) expressing the SARS-CoV-1 spike protein (Witte et al. 2023). The methods used were similar to those described previously (Ruiz et al. 2024), including generation of virus, infections and Illumina sequencing methods of virus output, which was harvested at 48 hours after the second passage. mAb C68.490 was tested at two concentrations, 50ng/ml and 250 ng/ml, against rVSV expressing SARS-CoV-1 (CUHK-W1).

## RESULTS

### Isolation of spike-specific monoclonal antibodies after re-infection

To examine the evolution of the antibody response in C68 after additional exposures to SARS-CoV-2 antigen compared to our first sampling (Guenthoer et al., 2024), we isolated spike-specific MBCs from peripheral blood mononuclear cells collected at one month after the second BTI (sample BTI-2), which was during a period when Omicron BA.5 was the dominant circulating variant. This individual previously experienced multiple SARS-CoV-2 antigen exposures, including a first BTI with the Delta variant one year prior and an intervening vaccination (WH-1) seven months prior to the second infection (Fig 1). MBCs that bound spike were isolated from the BTI-2 sample using SARS-CoV-2 Omicron XBB.1.5 and SARS-CoV-1 (CUHK-W1 strain) trimers (Fig 1). These two spike proteins were chosen to enrich for antibodies with broad activity against SARS-CoV-2 and/or SARS-CoV-1.

**Figure 1.**
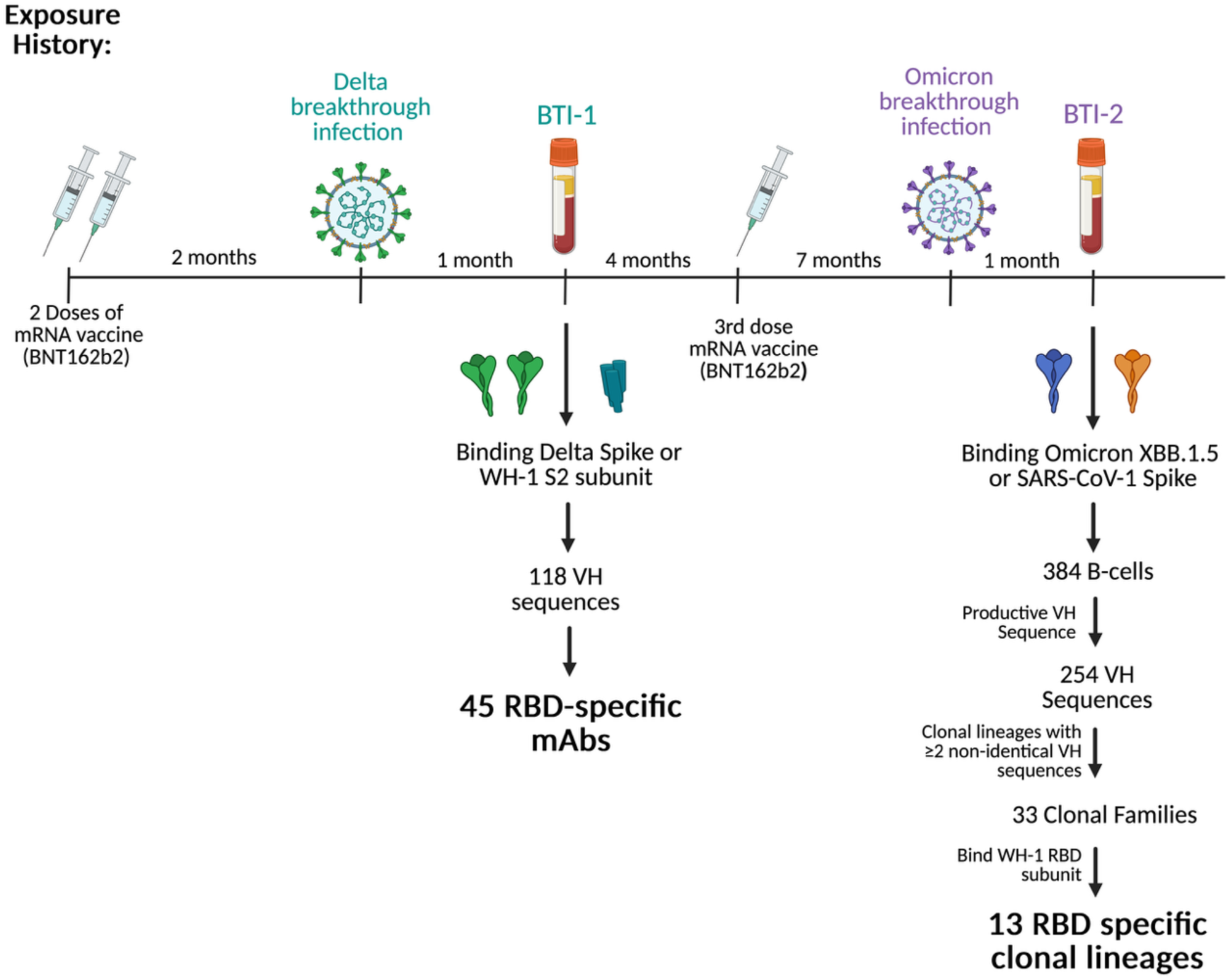
Schematic of individual C68’s exposure history with timing and approach for antibody isolation. Timeline indicating intervals of vaccination and infection is shown at the top. The time points at which antibodies were isolated (BTI-1 and BTI-2) are indicated. Below the timeline the antigens used to select antigen-specific memory B cells are shown. The process used to identify the 13 RBD-specific antibodies from BTI-2 used for downstream studies is shown to the right. (Created with Biorender.com)

To identify clonal lineages, we pooled VH sequences at the BTI-2 timepoint with those previously isolated from the BTI-1 timepoint one year prior. We identified 33 cases of expanded clonal lineages (Fig 1), which we defined as at least two non-identical VH members. Of those, 11 were clonal lineage families that were detected in both BTI-2 and BTI-1 samples and 22 were new clonal lineage families only found in the BTI-2 sample. Because most potently neutralizing antibodies target the RBD, this subset was further selected for those that bound to RBD by ELISA.

This resulted in four clonal lineages that include both BTI-1 and BTI-2 members and nine that were identified only in the BTI-2 sequences for downstream studies.

### BTI-2 antibodies show increased binding and neutralizing activity compared to clonally related BTI-1 antibodies

To address whether re-infection and associated continued antigen exposure impacts antibody breadth and function, we first focused on the four clonal lineages with antibodies from both BTI-1 and BTI-2. All four BTI-2 mAbs had higher levels of SHM than BTI-1 mAbs from the same clonal lineages based on VH chain sequences, but only clonal lineage 4 showed a notable change in SHM of 7.1% (from 3.1% to 10.2%). For the other three clonal lineages, increases in SHM were more modest, ranging from 0.2% to 1.1% (Fig. 2A). Changes in SHM were similarly modest for three of the VL sequences (ranging from an increase of 0.7% to 2.6%), with one lineage (clonal lineage 3) having a decrease in SHM from 4.6% to 2.9%.

**Figure 2.**
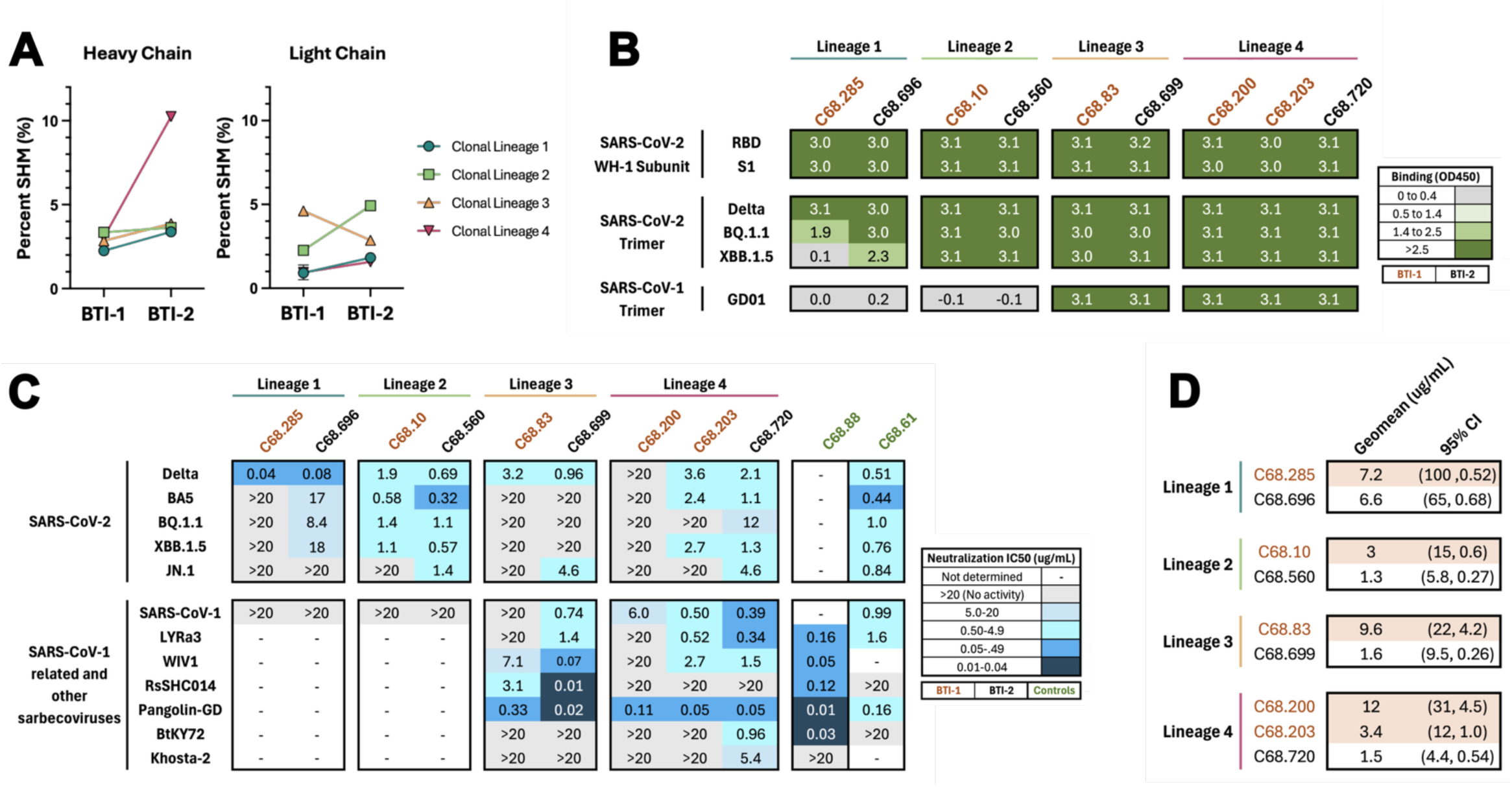
**Functional activity of BTI-2 antibodies compared to BTI-1 antibodies from the same clonal family.** A) Percent somatic hypermutation of IgG heavy chain and light chain among clonal lineage members. B) Comparison of binding profiles between BTI-1 mAbs and clonal lineage members in BTI-2. Heatmap shows OD450 nm binding activity by ELISA with the antigens tested shown to the left. The antibodies tested are indicated at the top with mAbs from BTI-1 in orange text and BTI-2 in black text. mAbs were tested in duplicate at a fixed concentration of 1000 ng/ml. Green indicates binding with increasing shades of green indicating increasing OD450 values, as indicated in the key to the right. Grey indicates no detectable binding. C) Heatmap of neutralization IC50s (ug/mL) of the same antibodies as in B. Viruses tested are shown to the left with SARS-CoV-2 variants in the top half and more diverse sarbecoviruses in the bottom half. The color gradient represents neutralization activity as shown in the key to the right, with darker shade of blue correlates with IC50 potency. Grey indicates no detectable neutralization (IC50) and white indicates the mAbs not tested because they did not bind SARS-CoV-1 spike trimer. IC50 values were averaged from 2–3 independent experiments performed in technical duplicate. IC50 values were calculated with GraphPad Prism, with a four-parameter non-linear regression model. C68.88 and C68.61 are positive control mAbs that were tested in parallel. D) Geometric mean of the IC50 values (ug/mL) and 95% Confidence Intervals (CI) of the panel of viruses tested in 2C. IC50 >20 ug/mL was set to 20 ug/mL this calculation.

The binding breadth of these mAbs was examined by ELISA against several spike proteins, including from the ancestral WH-1 strain as well as SARS-CoV-2 variants and SARS-CoV-1. All BTI-2 mAbs bound WH-1 RBD and S1 subunits, as well as Delta trimer (Fig 2B). One BTI-2 mAb, C68.696, showed increased binding breadth against Omicron variants relative to the BTI-1 mAb, C68.285, from the same clonal lineage (clonal lineage 1). In this case, C68.696 acquired the ability to bind the SARS-CoV-2 XBB.1.5 trimer compared to the related BTI-1 mAb. For the other three clonal lineages, the binding patterns were similar and for clonal lineages 3 and 4, both BTI-1 and BTI-2 mAbs bound all spike antigens tested.

We next compared the neutralization activity between these clonal mAbs against a small panel of SARS-CoV-2 variants, including Delta and Omicron variants (Fig. 2C). In all instances, BTI-2 mAbs showed increased neutralization (i.e., lower IC50) compared to their clonal lineage members from the BTI-1 time point. For example, the BTI-1 mAbs C68.285 from clonal lineage 1 neutralized only the Delta pseudovirus, whereas the BTI-2 member of this lineage, C68.696, acquired the ability to neutralize Omicron BA.5, XBB.1.5 and BQ.1.1. BTI-2 mAbs from all three other clonal lineages neutralized at least one virus more potently than their BTI-1 clonal counterparts.

The BTI-2 mAbs of clonal lineage families 3 and 4 neutralized SARS-CoV-1. For these clonal lineages, we further examined neutralization activity against a larger panel of sarbecoviruses. We pseudotyped viruses with spikes from representative viruses from three clades of sarbecoviruses. This included Pangolin-GD, a clade 1b virus closely related to SARS-CoV-2 (96% sequence identity to the ancestral SARS-CoV-2 RBD), and several clade 1a viruses related to SARS-CoV-1 including WIV1 (96% sequence identity to ancestral SARS-CoV-1 RBD), LYRa3 (95%), and RsSHC014 (82%). In addition, we tested two more distantly related viruses from clade 3 sarbecoviruses that include isolates from Africa and Europe, BtKY72 and Khosta-2 (73% and 68% to SARS-CoV-2 RBD, respectively). Because wildtype BtKY72 has reduced binding of human ACE2, we utilized a K493Y/T498W mutant that confers entry via human ACE2 (Starr, Zepeda, et al. 2022). We included two previously characterized BTI-1 mAbs that display breadth across SARS-CoV-2 variants (C68.61) or other sarbecoviruses (C68.88) and both showed the expected pattern of neutralization (Fig 2C).

In both clonal lineages, BTI-2 mAbs showed increased breadth and potency in neutralization against this panel of sarbecoviruses compared to BTI-1 mAbs. For instance, in clonal lineage 3, BTI-2 mAb C68.699 neutralized five of the seven sarbecoviruses tested compared to three of seven for the BTI-1 clonal lineage member (C68.83). These mAbs also showed differences in potency, for example, the BTI-2 mAb neutralized RsSHC014 with a 300-fold greater potency compared to its BTI-1 clonal lineage member (IC50 = 3.1 vs. 0.01 ug/mL). Similar increases in breadth and potency in sarbecovirus neutralization were also seen for clonal lineage 4. In this case, the two sarbecoviruses with the lowest sequence similarity to SARS-CoV-2, BtKY72 and Khosta-2, were not neutralized by either mAb from the BTI-1 timepoint, but both were neutralized by the BTI-2 mAb C68.720.

Overall, the BTI-1 mAbs from all four clonal lineages displayed lower geometric mean IC50s (Fig 2D) than the BTI-2 mAbs across the panel of SARS-CoV-2 and sarbecovirus variants. Thus, in each case, the neutralizing activity had increased for the BTI-2 mAbs when compared to those isolated a year prior at BTI-1, including against sarbecoviruses that were not used to select for antigen positive B cells or a known part of the patient’s exposure history.

### Characterization of antibodies representing BTI-2-specific clonal lineages

In addition to clonal lineage families represented in both BTI-1 and BTI-2, we also identified nine RBD-specific clonal lineages that were unique to BTI-2. For two lineages (6 and 13), two mAbs were included, resulting in a total of 11 RBD-specific antibodies. These clonal lineages utilized a diversity of VDJ genes. However, light chain gene usage was more homogenous, with five of nine clonal lineages utilizing IGVK1-5 (Supplemental Fig 2)

All 11 BTI-2 mAbs bound both SARS-CoV-2 WH-1 RBD and S1 subunits (Fig 3A). The majority bound XBB.1.5 and SARS-CoV-1 spike, with only 2 of 11 showing no or only weak binding to one or more spike variants. We assessed the ability of these mAbs to neutralize viruses pseudotyped with spikes from SARS-CoV-2 XBB.1.5 and SARS-CoV-1, which are the spike proteins used for isolation of MBCs from BTI-2. Most of the antibodies (8/11) had detectable neutralization (IC50 <20 ug/mL) against both XBB.1.5 and SARS-CoV-1 pseudoviruses, and all but C68.459 were capable of neutralizing at least one of these viruses. IC50s ranged from 1.3 to >20 ug/ml (mean IC50 = 10 ug/ml) against SARS-CoV-2 XBB.1.5. Interestingly, in all cases, except C68.459, these mAbs more potently neutralized SARS-CoV-1 than XBB.1.5 (Fig 3B), with IC50s ranging from 0.02 ug/ml to >20 ug/ml (average IC50 = 2.6 ug/ml). The most notable of these mAbs was C68.490, which neutralized SARS-CoV-1 at least 10-fold more potently than any other tested BTI-2 mAb (IC50 = 0.02 ug/mL). Furthermore, C68.490 was ∼50-fold more potent than C68.61 (IC50 = 0.99 ug/mL; Fig 2C), which we have previously reported as a potent RBD mAb isolated from BTI-1 that exhibits considerable sarbecovirus breadth (Guenthoer 2023, Ruiz 2024).

**Figure 3.**
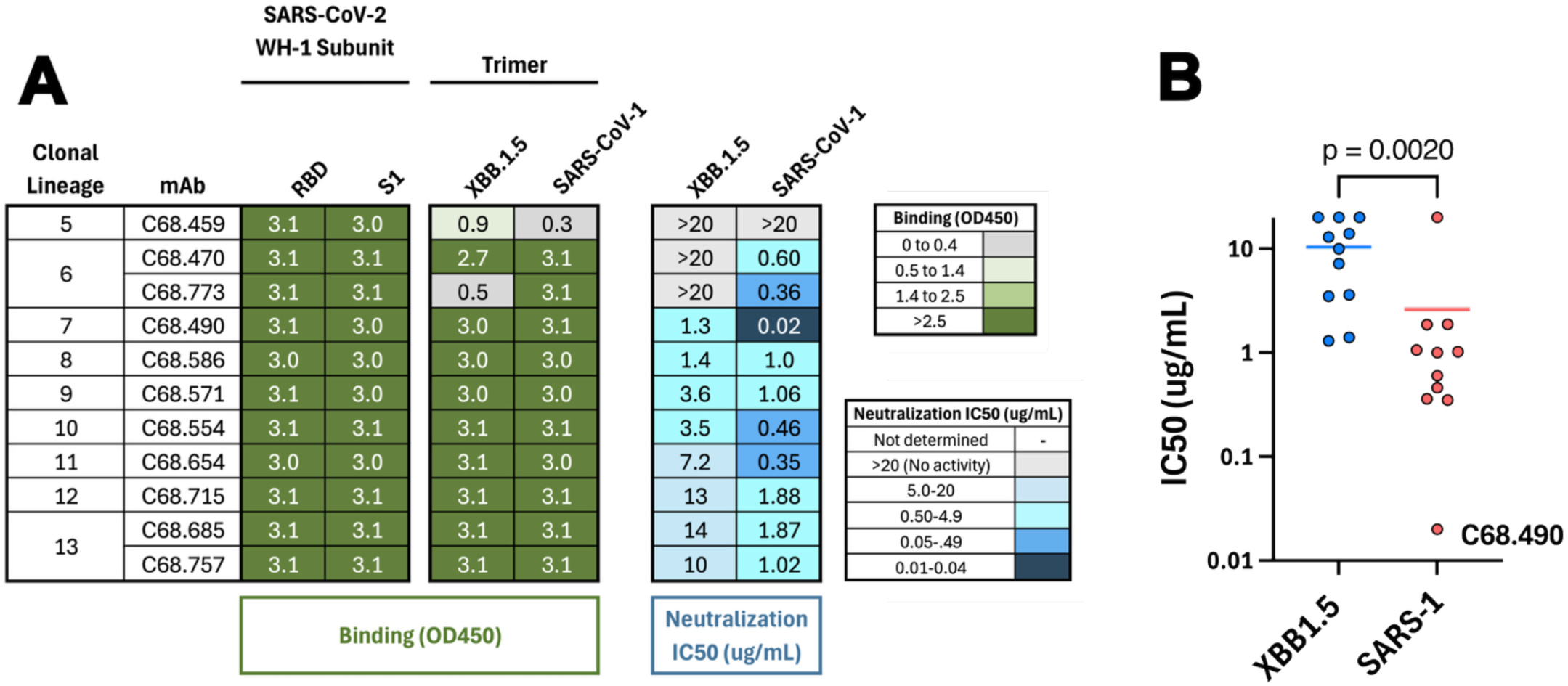
**Binding and neutralization of antibodies representing new clonal lineages specific to BTI-2.** A) Heatmap shows binding (green) and neutralization IC50s (blue, last two columns). Viruses tested are shown at the top and mAbs indicated to the left, after the clonal family designation. Figure details are as described in Figure 2B and 2C. B) Comparison of IC50s for SARS-CoV-2 XBB1.5 and SARS-CoV-1 for antibodies from BTI-2 specific clonal lineages. The p value is calculated from the Wilcoxon matched-paired signed rank test. Antibodies with no neutralization activity (IC50 >20 ug/mL) were set to 20 ug/mL for this comparison

### C68.490 demonstrates potent binding and neutralization across diverse sarbecovirus clades

Given that C68.490 showed potent neutralization activity against SARS-CoV-1 and also neutralized SARS-CoV-2 XBB.1.5, we sought to determine if this activity extended to other sarbecoviruses and additional SARS-CoV-2 variants. As a first step, we profiled binding breadth against a panel of 72 sarbecovirus RBDs using a yeast display system (Starr et al. 2021; Rosen et al. 2024). This library includes multiple variants within each sarbecovirus clade (clades 1a, 1b, 2 and 3) (Starr et al. 2021; Rosen et al. 2024) and thus includes RBDs from ACE2 receptor-utilizing sarbecoviruses, similar to SARS-CoV-1, SARS-CoV-2, and BtKY72, as well as RBDs from more divergent ACE2-independent viruses (clade 2). Remarkably, C68.490 bound the RBD of all 72 tested sarbecoviruses with EC50s less than 10 ng/mL, including potent binding to RBDs from the divergent non-ACE2-utilizing bat sarbecovirus clade (Fig 4A). Notably, the binding was broader than several of the most cross-reactive RBD mAbs previously described, including S2X259 (Tortorici et al. 2021; Cameroni et al. 2022), SA55 (Cao, Yisimayi, et al. 2022; Planas et al. 2024; Cao, Jian, et al. 2022) and VIR-7229 (Rosen et al. 2024) (Fig.4A). Among those mAbs, VIR-7229 showed the greatest breadth, with detectable binding to the RBD of all but a few members of the clade 1b family. C68.490 binding extended to all clade 1b RBDs and notably showed stronger binding to the clade 2 RBDs compared to VIR-7229 (Fig 4B); the other mAbs showed no (SA55) or limited (S2X259) detectable binding to RBDs from this clade. C68.490 binding was not statistically different from other antibodies against Clade 1b RBDs. Binding was slightly less compared to SA55 (mean EC50 = 0.24 vs. 0.20 ng/uL) for clade 1a RBDs and compared to VIR-7229 for clade 3 RBDs (mean EC50 = 1.2 vs 0.51 ng/uL).

**Figure 4.**
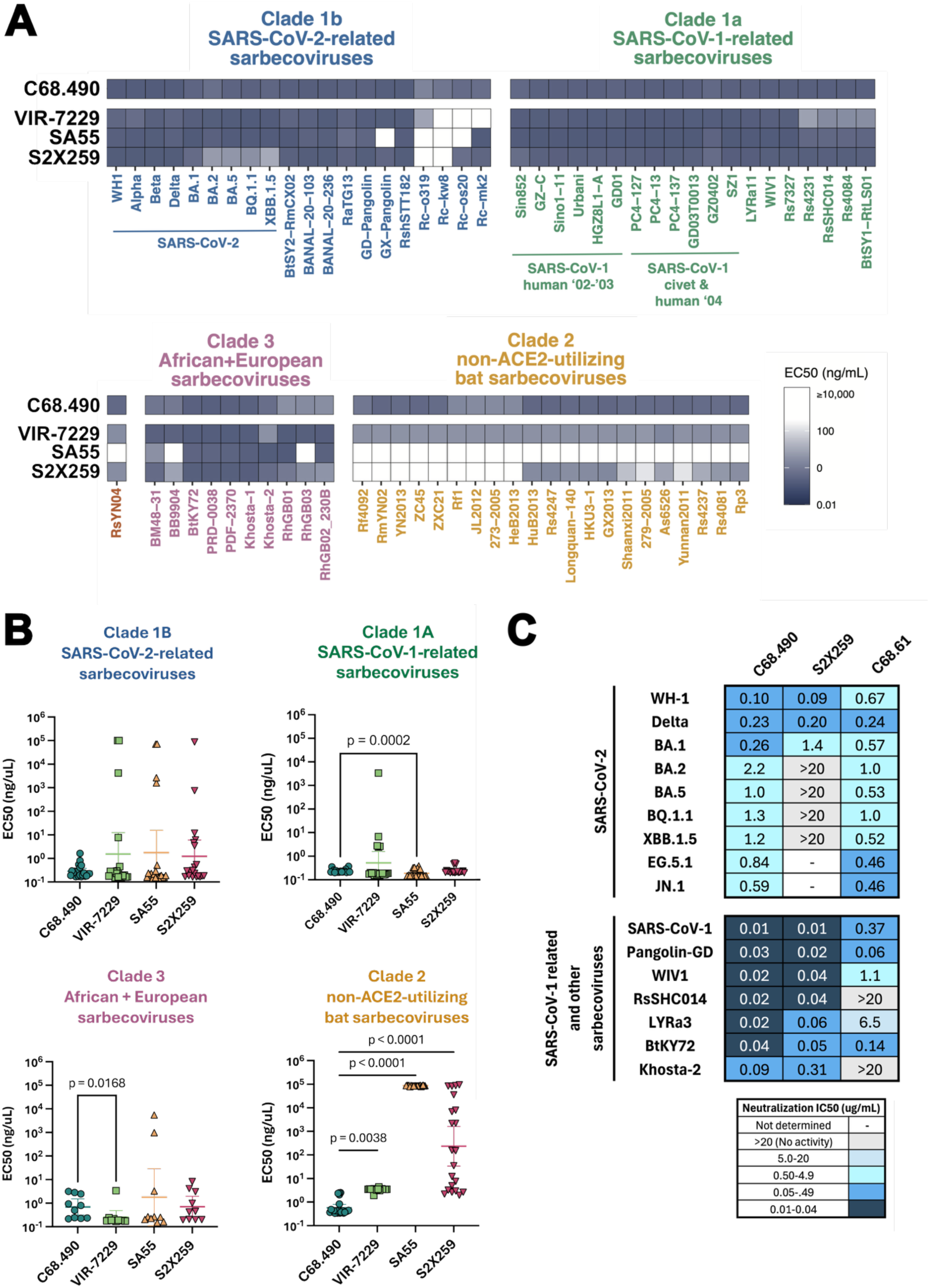
**Functional characterization of a pan-sarbecovirus neutralizing antibody C68.490.** A) Binding breadth of C68.490 and previously characterized antibodies with known pan-sarbecovirus activity against a library of yeast-display sarbecovirus RBDs. Antibodies tested are shown to the left with the specific RBD indicated at the bottom. RBDs are classified by clades as shown above and color-coded. Data for VIR-7229, SA55, and S2X259 previously published by (Rosen et al. 2024) and shown for comparison. B) Comparison of C68.490 binding with other pan-sarbecovirus antibodies by sarbecovirus clade. Statistically significant differences in mean EC50 values between C68.490 and other antibodies was assessed using the Friedman test with Dunn’s multiple comparisons test. Sarbecovirus RBDs were assigned to clades based on existing clade definitions (Starr, Greaney, et al. 2022). C) Neutralization of SARS-CoV-2 and sarbecovirus variants by C68.490. Antibodies tested are described at the top and include two control mAbs for comparison. The viruses tested are shown to the left, with a line separating SARS-CoV-2 variants from the more diverse sarbecoviruses. Neutralization data are represented as IC50s (ug/mL) and color-coded as shown in the table below. IC50 values where averaged from 2-4 independent experiments performed in technical duplicate, with the exception of neutralization of S2X259 against SARS-CoV-1, which was only tested once. Other details are as in Figure 2C.

To determine if C68.490 also demonstrated breadth in neutralization, we determined the neutralization IC50s of this antibody against major and recently circulating SARS-CoV-2 variants, as well as the ACE2-dependent sarbecoviruses described above. Among SARS-CoV-2 variants tested, C68.490 neutralized all viruses with IC50 measurements ranging from 0.10 ug/mL to 2.2 ug/mL (Fig 4C). While there was slightly reduced neutralization in BA.2 sublineage variants (IC50 = 0.59-2.2 ug/mL), in comparison to pre-BA.2 variants (IC50 0.1-0.26 ug/mL), C68.490 retained neutralization against more recently circulating variants EG.5.1 and JN.1 (IC50 = 0.84, 0.59), similar to C68.61, which was tested in parallel as an example of a mAb notable for its breadth against SARS-CoV-2 variants (Guenthoer et al. 2023; Ruiz et al. 2024).

C68.490 potently neutralized SARS-CoV-1 and other tested sarbecoviruses, with IC50s ranging between 0.01 to 0.09 ug/ml, including viruses with relatively low sequence similarity to SARS-CoV-2 like BtKY72 (IC50 = 0.04 ug/mL) and Khosta-2 (IC50 = 0.09 ug/mL). The breadth and potency of C68.490 against this sarbecovirus panel was similar to that of S2X259, which is one of the broadest cross-sarbecoviruses mAbs yet described (Tortorici et al. 2021). Thus overall, C68.490 is notable for its broad and cross-reactive neutralizing activity against both SARS-CoV-2 variants and diverse sarbecoviruses, consistent with its broad binding profile.

### C68.490 targets a highly conserved RBD epitope with limited escape pathways

To define the epitope of C68.490, we examined binding to a deep mutational scanning (DMS) library encoding virtually all single mutations within the RBD, allowing the identification of the functional epitope (Dingens et al. 2019) as sites where mutations escape antibody binding. Because the epitope profiling can be influenced by epistatic interactions with other amino acid residues (Starr, Greaney, et al. 2022; Witte et al. 2023), binding was assessed using DMS libraries in the background of two SARS-CoV-2 variants, WH-1 and BA.2 (Starr, Greaney, et al. 2022), as well as, for the first time, SARS-CoV-1 (Lee et al. 2023). For all three RBDs, the epitope of C68.490 was focused within sites 378-387, based on WH-1 numbering, with two shared binding escape sites, positions 378 and 384, across all three RBD proteins (Fig 5A). There were also unique escape sites in some backgrounds, such as mutations at 386 that led to reduced mAb binding for WH-1 and mutations at 383 and 387 that escape binding in SARS-CoV-1. A common feature of mutations that were most disruptive to binding across all three RBDs was a change to a negatively charged amino acid. At position 384, aspartic acid (D) was the most enriched amino acid escape mutation across all three RBDs.

**Figure 5.**
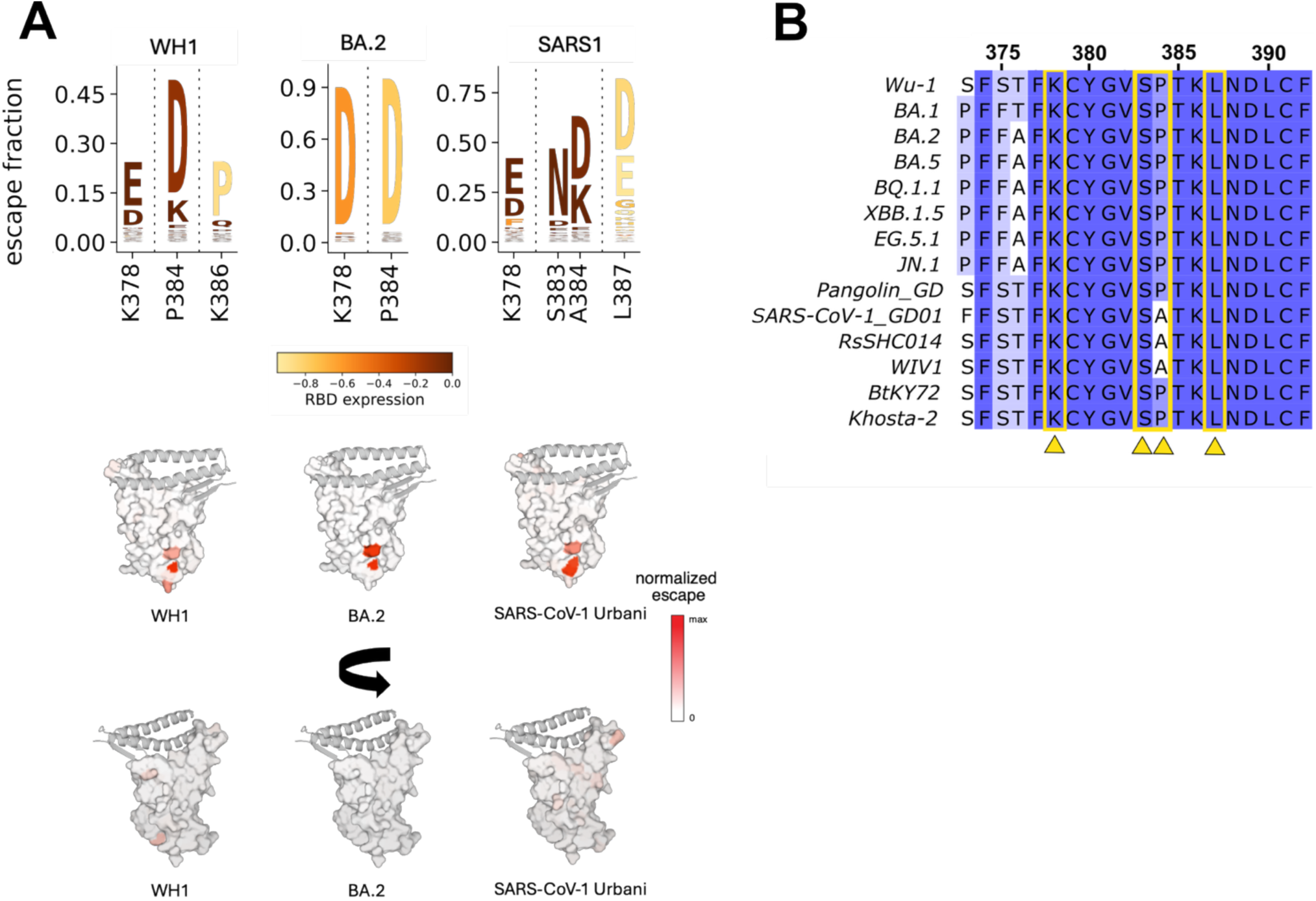
**Characterization of C68.490 escape mutations.** A) Top, Sites of binding escape and residues conferring escape from C68.490 in different viral backgrounds (WH1 = SARS-CoV-2 Wuhan-Hu-1, BA.2 = SARS-CoV-2 Omicron BA.2, SARS1 = SARS-CoV-1 Urbani) by deep mutational scanning of the RBD. Amino acid numbering based on SARS-CoV-2 WH1 sequence. Bottom, sites of escape mapped onto surface representation of the RBD (key interacting motifs of ACE2 shown as the gray ribbon). Color gradient represents escape fraction with darker red encompassing degree of escape. B) Multi-clade sarbecovirus sequence alignments. Conserved sites (in blue) and variable residues (in white) across sarbecoviruses around the epitope of C68.490 defined in A are depicted. C68.490 sites of escape are indicated with yellow arrows.

Several of the mutations that reduced binding of C68.490 to the RBD also reduce RBD expression, as defined in previous studies using these yeast RBD libraries to examine mutant effects on antibody escape and ACE2 binding (Starr, Greaney, et al. 2022; Lee et al. 2023; Starr et al. 2020) (Fig 5A, yellow color). This is most notable in the BA.2 background where both dominant escape mutants also reduce RBD expression, indicating that while these mutations reduce binding of C68.490, viruses encoding these may have impaired fitness due to poor RBD folding. Indeed, when we performed experiments using replication-competent recombinant VSV-G encoding SARS-CoV-1 CUHK-W1 spike we saw little selection for escape. At concentration 50 ng/mL, no escape was observed whereas at concentration 250ng/mL, there was only a minor fraction (14%) of the virus that had a mutation, and this was at a position, G381R, outside of the core epitope.

Sequence alignment of all escape sites identified in the DMS escape profiles illustrates that these sites are highly conserved across both SARS-CoV-2 variants as well as other sarbecoviruses sequences. Three out of four escape sites are completely conserved across all the viruses tested for neutralization (Fig 5B). At the fourth site, SARS-CoV-1, RsSHC014 and WIV1 encode an alanine at position 384 compared to a proline in the other RBDs. However, this amino acid substitution does not disrupt binding (Fig 5A), which is consistent with the finding that the presence of alanine 384 was not associated with a reduction in neutralization activity (Fig 4C).

In an analysis of 2910 geographically and temporally diverse SARS-CoV-2 genomes from Genbank at site K378 using Nextstrain (Hadfield et al. 2018), no mutations were observed (Supplemental Fig 3). At site P384, three mutations have been rarely observed. None of these mutations were predicted to lead to binding escape in any of the three viral backgrounds tested by DMS.

## DISCUSSION

Given the prevalence of sarbecoviruses in animal reservoirs and their demonstrated capacity to enter the human population, continued spillover events and associated pandemics are possible. The rapid emergence of variants during the SARS-CoV-2 pandemic illustrated that once the virus circulates in humans, there is rapid immune escape driven by mutations in the spike protein. Thus, to effectively target these viruses, antibodies must target conserved regions of the viral spike protein and have broad neutralizing activity. Here we built on studies suggesting that multiple exposures to diverse SARS-CoV-2 spike antigens can lead to broad plasma antibody responses (Wang et al. 2023, 2024; Abbad et al. 2024; Evans et al. 2022) by examining this response at the monoclonal level. mAbs isolated after re-infection showed increased breadth compared to those from the same clonal lineage isolated after the first infection with a different strain. Notably, the mAbs isolated after re-infection showed neutralizing activity against both SARS-CoV-2 and other sarbecoviruses, with one, C68.490, showing remarkable breadth against diverse sarbecoviruses. Thus, these mAbs, particularly C68.490, may be a valuable tool for prevention of or response to future sarbecovirus spillovers.

While studies of plasma antibody responses have shown that continued antigen exposure with diverse strains can lead to broader responses (Wang et al. 2023, 2024; Abbad et al. 2024; Evans et al. 2022), little was known about the details of this response. By comparing members of the same clonal lineage after a first breakthrough with the Delta variant to those present after a second breakthrough with an Omicron variant, we showed that some of this breadth reflects affinity maturation of antibodies within a lineage. This increased activity corresponded to a modest increase in SHM, a central part of the affinity maturation process. For two clonal lineages, the improved breadth was evident not only as expanded neutralization against SARS-CoV-2 variants, including Omicron variants, but also expanded activity against SARS-CoV-1 and related sarbecoviruses, which were notably not known to be a part of the C68 donor’s exposure history.

Several remarkable mAbs with broad activity against either SARS-CoV-2 or against sarbecoviruses from animal reservoirs have been reported (Cameroni et al. 2022; Tortorici et al. 2021; Pinto et al. 2020), but few have broad activity that encompasses both. C68.490 is thus unique in neutralizing across all of these viruses. Moreover, it shows high potency against sarbecovirus strains from different clades, including SARS-CoV-1. Remarkably, it bound to all 72 sarbecovirus spike RBD proteins tested, including more diverse non-ACE2 utilizing bat sarbecoviruses. Overall, the breadth of C68.490 exceeds that of previously described mAbs with notable activity across sarbecoviruses, including VIR-7229, SA55 and S2X259 (Rosen et al. 2024; Cao, Jian, et al. 2022; Cameroni et al. 2022) as well as C68.61, isolated from this individual after the first breakthrough infection (Guenthoer et al. 2023). These findings, along with results indicating that escape variants do not evolve under selection by C68.490 in culture, indicate that this mAb targets a highly conserved epitope present in diverse strains, suggesting functional constraints on this epitope.

The C68.490 epitope mapped to the core region of the RBD, including positions 378 and 384, which is a more conserved epitope that lends toward greater antibody breadth compared to mAbs that interact with the ACE2-binding surface of the RBD (Fan et al. 2022; Starr et al. 2021). Only a few mutations led to reduced binding of C68.490 to the SARS-CoV-2 spike, implying limited paths to escape. Mutations at positions 378 and 384 were found to reduce binding for both WH-1 and Omicron BA-2 strains, with additional selection at position 386 in the context of the ancestral WH-1 strain. Selected mutations to aspartic acid and glutamic acid were most common, suggesting that a change to an acidic amino acid may be critical for escape. Similarly, the mutations that enabled escape in the context of SARS-CoV-1 spike were primarily aspartic or glutamic acid. However, the escape profile for SARS-CoV-1 included four sites of escape: two that corresponded to those observed for SARS-CoV-2 and two others at nearby sites, 383 and 387. Notably, all of these sites are highly conserved across SARS-CoV-2 and sarbecovirus strains, with occasional mutations at position 384 that were not predicted to impact binding, suggesting C68.490 may bind an evolutionarily constrained epitope.

Monoclonal antibody therapy proved to be a valuable tool for reducing the severity of COVID-19 due to SARS-CoV-2 infection, especially before vaccines were available. Given that there have been multiple sarbecovirus spillover events in recent times, including SARS-CoV-2, which continues to cause significant morbidity and mortality, it is critical to continue to prepare for future introduction of new sarbecovirus infections in humans, leveraging what we have learned from SARS-CoV-2. Our studies suggest that repeated SARS-CoV-2 exposure with different strains may not only improve responses to that virus, but also elicit broad and potent responses to related viruses present in animal reservoirs. These responses, and the antibodies that drive them may provide valuable first line approaches to the next pandemic.

## Acknowledgments

This was supported by NIH grant AI138709 to JO and DP2 AI177890 and the Searle Scholars Foundation to TNS. DR was supported by U01 AI150747. FR was supported by an NSF training grant DGE-2140004 and DD by a Washington Research Foundation Fellowship. P.D.B. was supported by P01 AI165075. V.A.B was supported by the Boehringer Ingelheim PhD Fellowship. We thank Marceline Côté (University of Ottawa), Amit Sharma (Ohio State University), Jesse Bloom (Fred Hutchinson Cancer Center), David Veesler (University of Washington) and Pamela Bjorkman (Caltech) for generously providing spike plasmids for production of spike-pseudotyped lentiviruses. We also would like to thank the participants and the study staff of the Hospitalized or Ambulatory Adults with Respiratory Viral Infections (HAARVI) study.

