## Supplemental Figures for "Re-infection with SARS-CoV-2 is associated with increased antibody breadth and potency against diverse sarbecovirus strains"

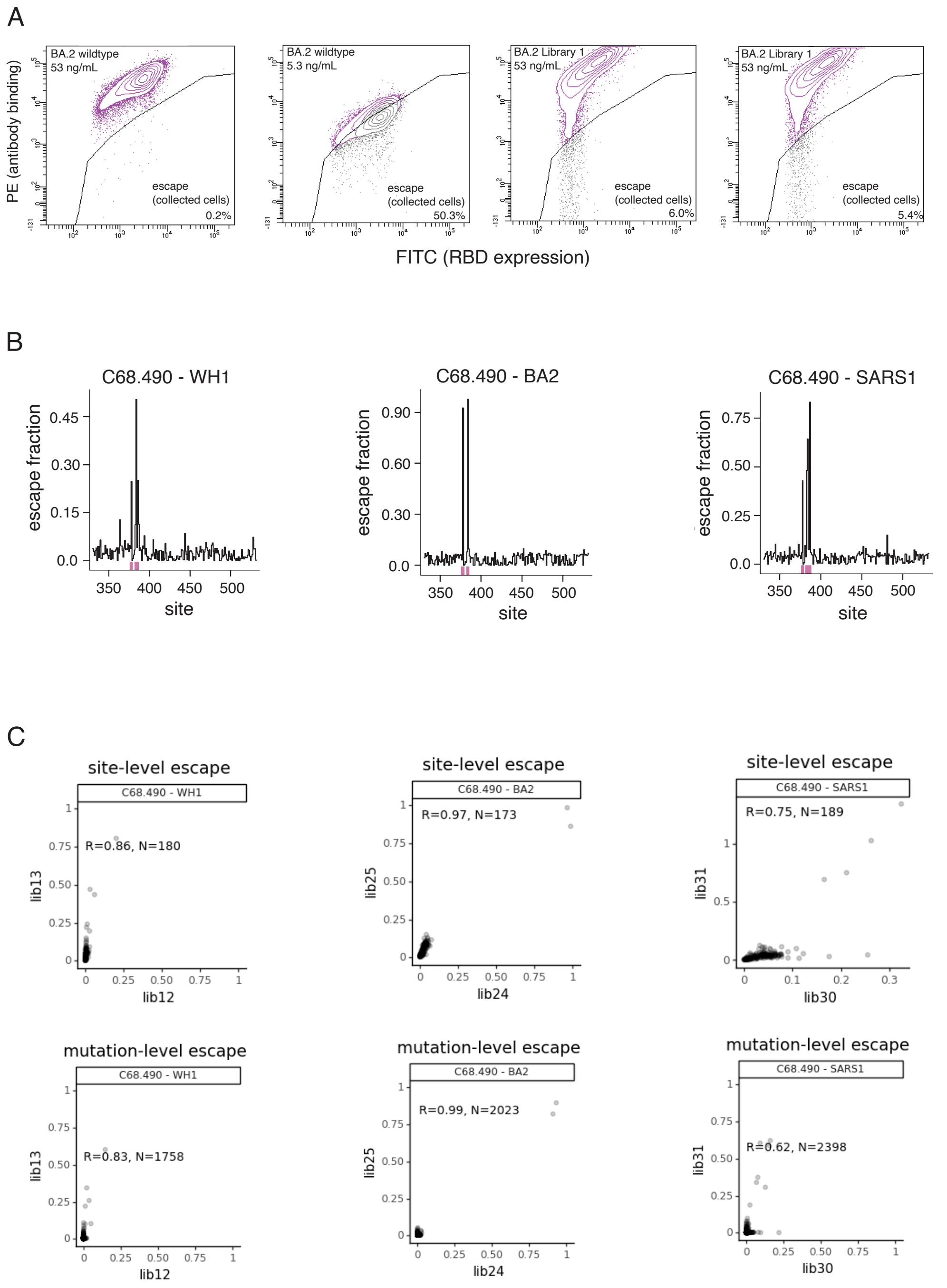


**Supplemental Figure 1. FACS Gating scheme, line plots of escape per site, and correlations of escape across replicates for yeast-display DMS escape selections.**

| | | | Heavy Chain | | | | | Light Chain | | | |
|---|---|---|---|---|---|---|---|---|---|---|---|
| **Clonal Family** | **mAb** | **Timepoint** | **V-gene** | **D-gene** | **J-gene** | **CDR3 Length** | **% SHM** | **V-gene** | **J-gene** | **CDR3 Length** | **% SHM** |
| 1 | C68.285 | BTI-1 | IGHV1-46 | IGHD5-24 | IGHJ4 | 13 | 2.3% | IGKV1-9 | IGKJ4 | 13 | 0.9% |
| | C68.696 | **BTI-2** | | | | | 3.4% | | | | 1.8% |
| 2 | C68.10 | BTI-1 | IGHV5-51 | IGHD3-3 | IGHJ4 | 14 | 3.4% | IGLV3-19 | IGLJ2 | 13 | 1.6% |
| | C68.560 | **BTI-2** | | | | | 3.6% | | | | 2.3% |
| 3 | C68.83 | BTI-1 | IGHV3-30 | IGHD2-15 | IGHJ6 | 24 | 2.8% | IGLV3-25 | IGLJ2 | 13 | 4.6% |
| | C68.699 | **BTI-2** | | | | | 3.9% | | | | 2.9% |
| 4 | C68.200 | BTI-1 | IGH3-23 | IGHD1-1 | IGHJ4 | 15 | 3.1% | IGKV1-5 | IGKJ5 | 10 | 0.6% |
| | C68.203 | BTI-1 | | | | | 3.1% | | | | 1.3% |
| | C68.720 | **BTI-2** | | | | | 10.3% | | | | 1.6% |
| 5 | C68.459 | **BTI-2** | IGHV4-31 | IGHD5-5 | IGH4 | 13 | 8.7% | IGKV4-1 | IGKJ2 | 11 | 3.2% |
| 6 | C68.470 | **BTI-2** | IGHV5-10-1 | IGHD4-17 | IGH4 | 17 | 5.5% | IGLV2-23 | IGLJ3 | 10 | 4.0% |
| | C68.773 | **BTI-2** | | | | | 4.6% | | | | 2.9% |
| 7 | C68.490 | **BTI-2** | IGHV1-18 | IGHD1-26 | IGHJ3 | 24 | 6.2% | IGKV1-5 | IGKJ4 | 11 | 3.4% |
| 8 | C68.586 | **BTI-2** | IGHV5-51 | IGHD2-21 | IGHJ4 | 13 | 7.2% | IGKV1-5 | IGKJ1 | 11 | 2.8% |
| 9 | C68.571 | **BTI-2** | IGHV4-59 | IGHD4-23 | IGHJ4 | 15 | 8.1% | IGKV1-33 | IGKJ4 | 11 | 5.0% |
| 10 | C68.554 | **BTI-2** | IGHV5-51 | IGHD2-21 | IGH4 | 15 | 7.8% | IGKV1-5 | IGKJ1 | 11 | 4.4% |
| 11 | C68.654 | **BTI-2** | IGHV3-64D | IGHD5-24 | IGHJ3 | 18 | 4.9% | IGKV1D-39 | IGKJ4 | 10 | 1.6% |
| 12 | C68.715 | **BTI-2** | IGH3-23 | IGHD1-1 | IGH4 | 15 | 6.1% | IGKV1-5 | IGKJ1 | 11 | 0.6% |
| 13 | C68.685 | **BTI-2** | IGH3-23 | IGHD2-15 | IGH4 | 15 | 7.5% | IGHK1-5 | IGKJ2 | 11 | 1.9% |
| | C68.757 | **BTI-2** | | | | | 8.9% | | | | 3.1% |

**Supplemental Figure 2. Gene Family usage for all RBD clonal antibodies**

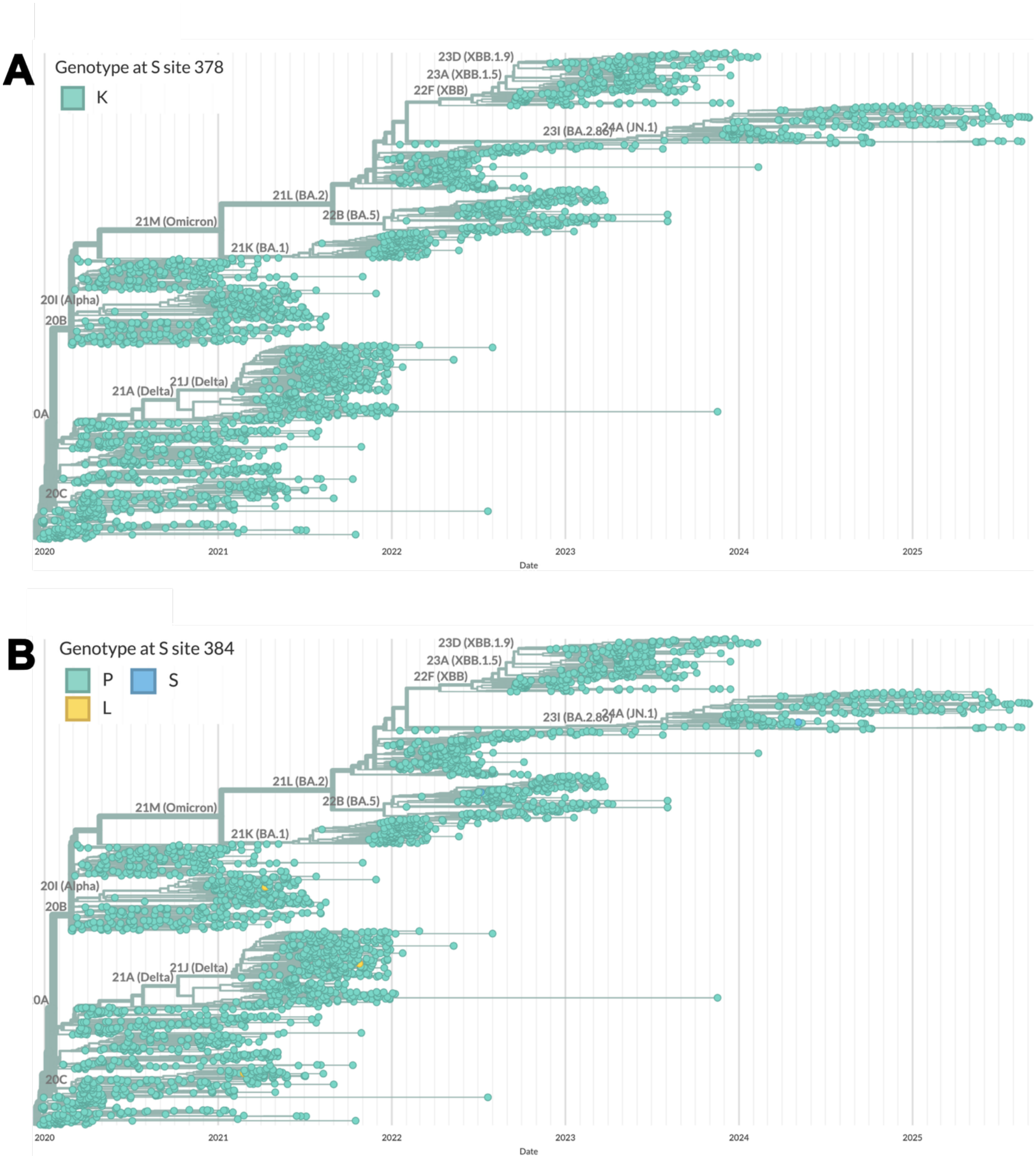


**Supplemental Figure 3. Nextstrain analysis of C68.490 key escape sites**

Genotype at SARS-CoV-2 spike amino acid sites 378 (A) and 384 (B) from 2910 genomes GenBank sequences (Retrieved Sept. 23, 2025 from Nextstrain.org).
